# Evidence for adaptive myelination of subcortical shortcuts for visual motion perception in healthy adults

**DOI:** 10.1101/2023.01.11.523655

**Authors:** Elise G. Rowe, Yubing Zhang, Marta I. Garrido

## Abstract

Conscious visual motion information follows a cortical pathway from the retina to the lateral geniculate nucleus (LGN) and on to the primary visual cortex (V1) before arriving at the middle temporal visual area (MT/V5). Alternative subcortical pathways that bypass V1 are thought to convey unconscious visual information. One flows from the retina to the pulvinar (PUL) and on to MT; while the other directly connects the LGN to MT. Evidence for these pathways comes from non-human primates and modest-sized studies in humans with brain lesions. Thus, the aim of the current study was to reconstruct these pathways in a large sample of neurotypical individuals and to determine the degree to which these pathways are myelinated, suggesting information flow is rapid. We used the publicly available 7T (N = 98; ‘discovery’) and 3T (N = 381; ‘validation’) diffusion MRI datasets from the Human Connectome Project to reconstruct the PUL-MT and LGN-MT pathways. We found more fibre tracts with greater density in the left hemisphere. Although the left PUL-MT path was denser, the bilateral LGN-MT tracts were more heavily myelinated, suggesting faster signal transduction. We suggest that this apparent discrepancy may be due to ‘adaptive myelination’ caused by more frequent use of the LGN-MT pathway that leads to greater myelination and faster overall signal transmission.

## INTRODUCTION

The human brain contains more than 80 billion neurons that transmit information rapidly via a combination of cortical and subcortical pathways (Rhoton, 2019). While each pathway might subserve a different underlying function, mounting evidence suggests that many carry redundant information (McFadyen et al., 2020). Surprisingly, little is known about the redundancy and efficiency of many of these pathways that support vital functions and survival-based behaviours. Of the visual signals required for survival-based threat detection, fear signalling and motion perception are two of the most important (Carr, 2015; Periera & Moita, 2016). Fear triggering is required to initiate the ‘fight or flight (or freeze)’ response to impending threat (Cannon, 1915; Phelps & LeDoux, 2005), while motion detection is necessary both to alert the organism to an approaching predator and to guide escape if the flight response is activated (Mobbs et al., 2015; Ferreira & Moita, 2020). Fear responses in humans are modulated by the amygdala via a well-studied cortical pathway (Pessoa & Adolphs, 2011) and a recently confirmed subcortical pathway that passes information directly from the retina to the superior colliculus then on to the pulvinar and amygdala (McFadyen et al, 2019; Koller et al., 2019). The benefit of the redundancy of this dual-route lies in the ability of the longer (and slower) *cortical* pathways to carry more detailed sensory information while the shorter *subcortical* route can rapidly transmit coarse sensory signals required for reflexive survival-based behaviours (Le Doux, 2000; Garrido, 2012). Thus, these subcortical pathways represent an efficiency mechanism to rapidly transmit information most important for survival. It is unclear, however, whether such subcortical shortcuts also hasten vital signals relating to visual motion.

Visual motion information follows a cortical pathway from the retina to the lateral geniculate nucleus (LGN; the visual thalamic nuclei) and on to V1 before arriving at the middle temporal visual motion area (MT/V5; Albright, 1984; Clifford et al., 2007). While V1 neurons represent position, size, orientation, motion and colour information across the visual field (Tootell et al., 1988), neurons in area MT specialise further, selectively representing the direction and speed of stimulus motion (Albright, 1984). Importantly, the neural activity in MT is directly linked to the perception of motion as measured by behaviour using non-human primates (Britten et al., 1992; Salzman et al., 1992). First, Britten et al. (1992), determined that perceptual performance matches the neural activity in MT, and Salzman et al. (1992), showed how this perceptual performance could be modified by electrically stimulating MT neurons. Area MT is reciprocally connected with V1 - V4, the middle superior temporal region and the ventral intraparietal area (Maunsell & Van Essen, 1983). Together, these regions form the major structures of the dorsal (cortical) visual processing pathway; with each region having distinct processing specificities relating to visual motion (Rizzolatti & Matelli, 2003). The subcortical pathways involved in motion processing are currently under debate, however, two thalamo-cortical pathways have been identified based on data from non-human primates and in ‘blindsight’ patients with V1 damage.

These two shortcuts flow from the retina to the pulvinar (PUL) and on to MT (for review see Bourne & Morrone, 2017), and from the LGN (which also receives direct input from the retina) directly to MT, bypassing V1 entirely (see Bridge et al., 2008 and for review see Tamietto & Morrone, 2016). Evidence for the PUL-MT pathway comes from the neonatal developmental physiology of marmosets in which strong, direct connectivity between the medial division of the inferior pulvinar and MT is observed (see Bridge et al., 2016). This pathway regresses during normal development to adulthood as the LGN-V1 pathway encourages the dominance of the optic radiations to V1, and then on to MT (Warner et al., 2010; Warner et al., 2015; Mitchell and Leopold, 2015). Thus, it has been suggested that the early maturation of MT may rely on early-life influences from the PUL (Warner et al., 2015). Conversely, evidence for the second subcortical pathway between LGN-MT comes primarily from blindsight patients who, after destruction of parts of V1, retain remarkable abilities to detect changes within their ‘blind field’ for properties associated with the dorsal visual pathways (e.g., motion and luminance contrast; Bridge et al., 2008). This pathway has been suggested as underlying blindsight capabilities in both monkeys (Sinchich et al., 2004; Schmid et al., 2010; Yu et al., 2018) and humans (Bridge et al., 2008; Ajina et al., 2015; ffytche et al., 1995; Ganglianese et al., 2012). However, it should be noted that a handful of studies suggest that the PUL-MT pathway might also contribute to some blindsight capabilities as evidenced in monkeys (using pharmacological inactivation of PUL, Kinoshita et al., 2019) and humans (observed in one patient with a gestational tumour within V1; Aghakhanyan et al., 2004). Critically, one of the major limitations of these prior studies, as outlined above, is that evidence for both pathways comes from non-human primates, or from patients with atypical neurodevelopment and/or neuroanatomy (and age-matched controls) in studies with small sample sizes.

Accordingly, the aim of the current study was threefold: (1) to reconstruct these two subcortical pathways between PUL-MT and LGN-MT in a large sample of neurotypical individuals, (2) to examine which regions are more heavily connected, as evidenced by a higher number of streamlines and greater fibre densities, and (3) to determine the extent to which these pathways are myelinated, suggesting neural signals are transmitted more rapidly (Caminiti et al., 2012; Chopra et al., 2018). To this end, we utilised the publicly available MRI and diffusion tensor imaging (DTI) datasets from the Human Connectome Project. We used both the 7T (N = 98) and 3T (N = 381) imaging resolutions as ‘discovery’ and ‘validation’ datasets, respectively. The ‘discovery’ dataset was used to examine our primary hypotheses, while the ‘validation’ dataset was used to confirm these results. We traced the two pathways proposed to underlie rapid motion perception between PUL-MT and LGN-MT using diffusion tensor imaging (DTI) fibre tractography. We used the T1w/T2w ratio as a proxy for myelin content (Ganzetti et al., 2014). The results of this study are important for identifying the brain pathways underlying visual motion processing that potentially enable rapid detection of change and threat.

## METHODS

### Participants

We utilised the 7T and 3T publicly available Human Connectome Project (HCP; van Essen et al., 2013) datasets as discovery (7T) and validation (3T) sets to strengthen the validity of our findings. The HCP 7T release contained data from 184 healthy young adults (mean age = 29.4 +/− 3 years). Of these, we excluded those who tested positive for drugs (alcohol, cannabis, meth/amphetamines, cocaine and opioids) and then randomly selected a subsample of N = 98 (mean age = 29.4 +/− 3 years) that did not contain any twins or non-twin siblings. The HCP 3T release contained the data from 1,206 healthy young adults (mean age = 28.8 +/− 4 years). Of these, we again excluded those testing positive for a number of drugs and then randomly selected a subsample of N = 381 (mean age = 28.7, +/− 4 years) excluding siblings. All participants in both datasets had normal or corrected-to-normal vision and no history of significant neurodevelopmental, neurological or neuropsychiatric disorders.

### dMRI acquisition - 7T

The HCP 7T dataset was collected in a Siemens Magnetom 7T Magnetic Resonance Imaging (MRI) scanner, a Nova32 32-channel Siemens head coil and a head-only transmit coil that surrounded the receiver head coil. Diffusion MR images (dMRI) were acquired over four sessions (each around 9.5 minutes) representing 2 different gradient tables (each with both anterior-to-posterior and posterior-to-anterior phase encoding polarities). Each gradient table included approximately 65 diffusion weighting directions (plus six b=0 acquisitions randomly collected throughout each run) and two shells with b-values of 1000 and 2000 s/mm^2^. Further parameters included: voxel size = 1.05 mm isotropic, TR = 7000 ms, TE = 71.2 ms, flip angle = 90 degrees, refocusing flip angle = 180 degrees, FOV = 210×210 (RO × PE), slice thickness = 1.05 × 1.05 mm and 132 slices, echo spacing = 0.82 ms, bandwidth = 1388 Hz/pixel.

### dMRI acquisition - 3T

The HCP 3T dataset was collected using a Siemens 3T ‘Connectome Skyra’ with a 32-channel head coil. Diffusion MR images were acquired in a single session with the following parameters (van Essen et al., 2013): three shells with b-values of 1000, 2000, and 3000 s/mm^2^; 90 directions per shell; voxel size = 1.25 mm isotropic; field of view = 210 × 180 mm; matrix = 168 × 144; slice thickness = 111 slices × 1.25 mm; TR = 5520 ms; TE = 89.5 ms; flip angle = 78 degrees; refocusing flip angle = 160 degrees; bandwidth = 1488 Hz/pixel; echo spacing = 0.78 ms.

### Structural MRI acquisition

The same 3T protocol was used for structural MRI acquisition, for both the 3T and 7T acquisition. T1- and T2-weighted whole-brain gradient-echo echo planar structural imaging (EPI) data were acquired at 3T (0.7 mm isotropic resolution) using a Siemens 3T ‘Connectome Skyra’ with a 32-channel head coil and a scan time of about 8 minutes. The T1-weighted images used a 3D MPRAGE sequence with TR = 2,400 ms, TE = 2.14 ms, TI = 1,000 ms, flip angle of 8 degrees, FOV = 224 × 224 mm and bandwidth = 210 Hz/pixel. The T2-weighted images used a 3D T2-SPACE sequence with TR = 3,200 ms, TE = 565 ms, a variable flip length, FOV = 224 × 224 mm and bandwidth = 744 Hz/pixel. To account for the use of the 3T structural images with the 7T diffusion data, the 3T images were collected at a resolution of 1.6 mm with a 59k mesh surface. Two separate volumes were collected for both the T1- and T2-weighted images and we used the first of these two scans in our analyses.

### dMRI pre-processing

We used the minimally processed 7T and 3T diffusion images provided by the HCP (van Essen et al., 2013). The minimal preprocessing pipeline included corrections for spin-echo-fieldmap susceptibility, eddy current, and motion distortions. These images were accompanied by motion parameters across which the mean relative displacement between dMRI volumes could be calculated. All further analyses were conducted using MRTrix 3.0.3 (Tournier et al., 2012), FSL 1.3.0 (Jenkinson et al., 2012) and SPM12 (http://www.fil.ion.ucl.ac.uk/spm/).

### Response function and white-matter fibre-orientation estimation

We generated response function estimates by segmenting each participants’ DWI image into the three tissue types of white matter, grey matter, and cerebrospinal fluid using the Dhollander et al., (2016b) algorithm. This algorithm provides unsupervised estimation of the tissue-types without requiring the subject’s T1 image and improves upon previous methods of obtaining single-fibre white-matter response function estimates (Dhollander et al., 2019). Next, to enable later-stage normalisation across the participant group, we estimated the average (or representative) response functions for each of the three tissue-types using the data from all individuals (separately for the 7T and 3T participant groups). We then used Multi-Shell, Multi-Tissue (MSMT) Constrained Spherical Deconvolution (CSD; Jeurisson et al., 2014; Dhollander et al., 2016a; Dhollander et al., 2019) to obtain the participant’s white-matter (WM) fibre-orientation distribution (FOD) from the underlying DWI data.

### Global intensity normalisation

Finally, to account for global intensity differences between participants and allow for comparison of the white matter tracts across the group, we applied a normalisation measure to individual participants WM FODs by inputting the average tissue response function components across the group and correcting for any residual intensity inhomogeneities. The resulting normalised WM FODs could then be used for tractography analysis with meaningful comparisons across participants.

### ROI analysis

We were interested in examining the white matter pathways connecting the PUL-MT and LGN-MT. As such, we selected these regions as our ROIs in both hemispheres. We created masks of these ROIs in standard MNI space using FSL (Smith et al., 2004; Jenkinson et al., 2012). For the MT and LGN masks, we used the probabilistic Juelich Histological Atlas (Amunts et al., 2005) at a threshold of at least 50% probability. For the PUL ROI, we used the mask provided by McFadyen and colleagues (2019), which were based on the five subcompartments of the PUL (that is, the superior, inferior, anterior, medial and lateral PUL) identified using the same Juelich Atlas and in which any holes between the clusters were filled manually using FSL. Note that to further investigate the connections of the PUL in later stages of the analysis, we also included individual masks for each of the 5 known subcompartments. All masks were warped into standard MNI space and then to each participant’s DTI space, and we ensured there was no overlap between the PUL and LGN ROIs.

### dMRI analysis

For our tractography analyses, we selected two different white matter reconstruction methods as a means of cross-method validation: global and local probabilistic tractography. The global probabilistic tractography method used a Bayesian approach to reconstruct the whole-brain fibre configurations using a generative model to best explain the data across the entire brain. Here, the white matter tracts between our ROIs (LGN-MT and PUL-MT) were identified by examining the entire brain and selecting only the tracks that originated and terminated within two ROIs (LGN and MT or PUL and MT, and did not enter the alternative excluded ROI, PUL or LGN, respectively). The local probabilistic tractography method also used a generative Bayesian modelling approach based only within the ROIs (i.e., we did not generate a tractogram for the entire brain) to determine tracts that originated and terminated in a single direction based on the underlying white matter FODs (Tournier et al., 2010). Using both the global and local tractography approaches, we estimated the ipsilateral white matter tracts between the PUL-MT and LGN-MT within each hemisphere and did not examine any tracts that crossed the corpus callosum. Note that for either pathway, we also supplied the alternative ROI as an exclusion zone, meaning that any PUL-MT tract that entered the LGN or any LGN-MT tract that entered the PUL was excluded.

### Global tractography

To estimate the whole-brain tractography, we used a 50 million streamline estimate of the white matter tracts and reduced this down to 10 million tracks (representing a more biologically accurate measure of the apparent fibre density) using Spherical-Deconvolution Informed FIltering of Tractograms (SIFT) which filtered the whole-brain tractogram by weighting each streamline by a cross-sectional multiplier directly related to the underlying data (Smith et al., 2013, Tournier et al., 2019). In applying SIFT, we were accounting for the fact that longer or more pronounced white matter pathways tend to be overestimated due to their prominence which subsequently results in a greater number of ‘hits’ during probabilistic tractography reconstruction. That is, the SIFT algorithm filters the tractograms based on a proportional reconstruction of fibre populations across the whole-brain. After applying these methods, we then examined the white-matter pathways connecting our ROIs in either hemisphere.

### Local tractography

We used local probabilistic tractography as a less conservative method to examine our primary research question. For this, we used the iFOD2 algorithm (Tournier et al., 2010) and planted 10,000 unidirectional seeds originating in each of our ROIs. Thus, we obtained tractography estimates in the forwards and backwards directions between: (1) PUL-MT and (2) LGN-MT, in either hemisphere. Next, we applied SIFT2 (i.e., SIFT version 2; Smith et al., 2015; Tournier et al., 2019) to enable investigation of the apparent fibre densities of each track using the crosssectional multiplier weights representing the proportional weighting of each tract connecting each ROI. For visualisation purposes, we took each participant’s tractography results, warped them into standard MNI space and combined them for the group-level figures.

### Statistics: Tract apparent fibre density, count and length

To quantify the existence of each tract across participants, we first examined the proportion of participants for whom each tract was present (i.e., at least one fibre connecting the ROIs) using both the global and local tractography methods. Within this group of participants, we then determined the streamline count and apparent fibre density (AFD) values that represented the number of streamlines within each pathway. Two-way ANOVAs with the Main Effects of region (i.e. PUL or LGN) and hemisphere (i.e. left or right) were then used to examine any significant differences.

### Myelin Maps

Once the global and local tractography were complete, we wanted to determine the mean myelination along the pathways connecting the ROIs. As a proxy for the whole-brain myelin maps of each participant, we used the T1w/T2w ratio as described by Ganzetti and colleagues (2014). For this, we used the SPM add-on MRTool toolbox and input the T1w and Tw2 structural images for each participant. We then implemented bias correction and calibration to normalise the images before estimating the T1w/Tw2 ratio images. Once computed, we converted the tractography results of each participant into individual masks and extracted the mean myelination (or T1w/T2w ratio) along the PUL-MT and LGN-MT pathways for each participant. Note that we examined the tracts within hemispheres; meaning that for the local results we combined the tracts in both the forwards and backwards directions.

### Statistics: Mean myelination per tract across participants

Once we had obtained each participant’s myelination maps and reduced these to only the voxels along their individual pathways, we extracted the myelination values (i.e. the T1w/T2w ratio) across all participants. We quantified the mean myelination along each pathway across participants by calculating the average T1w/T2w value along each tract within each hemisphere. Two-way ANOVAs with the Main Effects of region (i.e. PUL-MT or LGN-MT) and hemisphere (i.e. left or right) were then used to examine any significant differences in the mean myelination values along the paths. We visualise the group-mean myelination map by warping the tracts (and corresponding T1w/T2w values) into standard MNI space, combining across participants and then taking the average.

## RESULTS

We used DTI-based probabilistic white-matter tractography to reconstruct two subcortical visual motion pathways: (1) PUL-MT and (2) LGN-MT. We then determined which pathway was more highly myelinated and, thus, could be inferred to transmit neural signals more rapidly. For this, we utilised both the 7T (N = 98) and 3T (N = 381) dMRI datasets made publicly available via the HCP (Van Essen et al., 2013; note that we removed all twin and non-twin siblings from the total participant pools, as well as people who tested positive for a number of drugs). We used the 7T data as our ‘discovery’ dataset (to investigate our primary research question) and then applied the same methods to the 3T data as our ‘validation’ dataset (to support any conclusions drawn from the higher resolution 7T results). The first step to address our primary research questions was to reconstruct the ipsilateral white matter pathways between our ROIs of interest: (1) PUL-MT and (2) LGN-MT using two complementary tractography methods: global and local tractography.

### Evidence for two subcortical routes carrying visual motion information

#### Global tractography

In our global tractography analyses we first reconstructed the whole-brain white matter structures based on each individual’s diffusion weighted images using a probabilistic Bayesian approach that first generated a total of 50 million white matter tracts before reducing these down to 10 million using SIFT (Smith et al., 2013). We then restricted our analyses to only the fibre pathways connecting (i.e., either originating or terminating in) the ROIs of PUL and MT (PUL-MT pathway, excluding tracts passing through the LGN) or LGN and MT (LGN-MT pathway excluding tracts via the PUL). Using this tractography method, we were able to reconstruct both pathways in healthy individuals in both the discovery and validation datasets. In the 7T discovery dataset, we found that the percentage of individuals having at least one fibre connecting the ROIs was 60% for the left PUL-MT, 70% for the right PUL-MT, 32% for the left LGN-MT and 41% for the right LGN-MT. In the 3T validation set, these reduced to 34% for the left PUL-MT, 31% for the right PUL-MT, 9% for the left LGN-MT and 11% for the right LGN-MT.

Based on the poor performance of the global tractography algorithm in reconstructing these long-range subcortical to cortical pathways we opted to focus on local probabilistic tractography instead. The primary difference between the two methods being that while global tractography creates a whole-brain tractogram and filters out the tracts between the ROIs, local tractography instead uses the underlying white matter fibre orientations (i.e., the WM FODs) and an iterative probabilistic algorithm to generate streamlines that seed and terminate only within the ROIs in either direction. As such, local tractography is less conservative but also less restricted by computational processing capacities (Tournier et al., 2019; Smith et al., 2012).

#### Local tractography

Using local probabilistic tractography, we observed evidence for ipsilateral pathways connecting both the PUL-MT (Figure 1A) and the LGN-MT (Figure 1B) in both hemispheres and in the majority of individuals. In the 7T discovery dataset, the percentage of individuals having at least one fibre connecting the ROIs for the left and right PUL-MT was 94% and 87%, respectively, and the LGN-MT 97% and 91%, respectively. In the 3T validation dataset, the proportion of individuals with at least one tract for the left and right PUL-MT was 97% and 91%, respectively, and the LGN-MT 95% and 89%, respectively.

**Figure 1.**
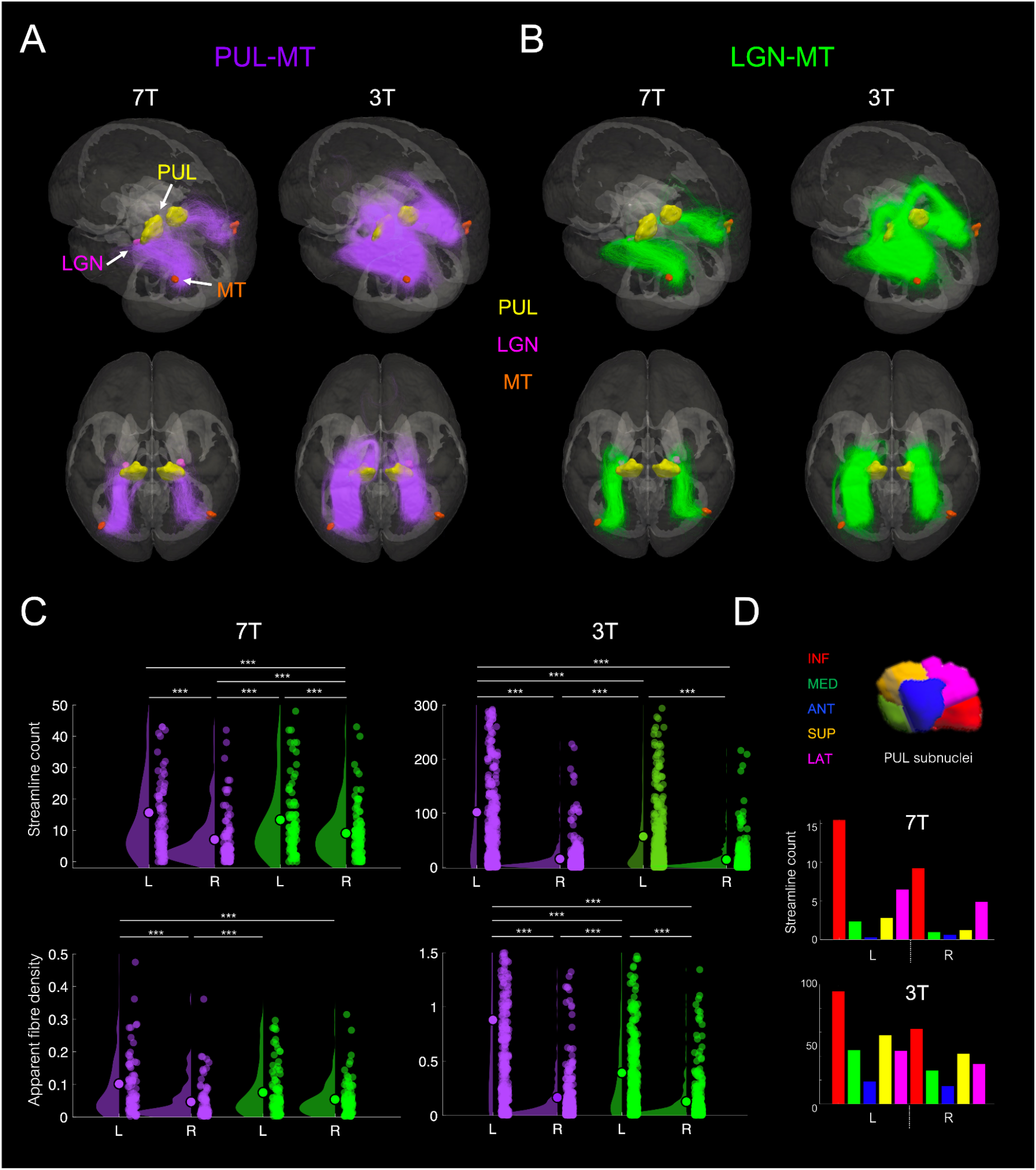
Local probabilistic tractography reconstruction of the white matter pathways connecting the pulvinar (PUL) and lateral geniculate nucleus (LGN) to visual area MT. **(A)** White matter pathways between the PUL-MT for the 7T ‘discovery’ dataset (left) and 3T ‘validation’ dataset (right). The top and bottom rows show the same data from a different perspective. **(B)** White matter pathways between the LGN-MT for the 7T ‘discovery’ dataset (left) and 3T ‘validation’ dataset (right). **(C)** Streamline counts (top) and apparent fibre density (bottom) for each participant within the left and right PUL-MT (purple) and LGN-MT (green) pathways for both 7T (left) and 3T (right) datasets. **(D)** Average number of streamlines terminating in each of the PUL subnuclei for 7T (top) and 3T (bottom) datasets. INF = inferior, MED = medial, ANT = anterior, SUP = superior and LAT = lateral.

### More tracts in the left hemisphere with greater density

#### Fibre counts

Next, we examined the number of streamlines (or white-matter fibre bundles) connecting the ROIs for each individual to determine which ROIs were more connected (Figure 1C, top). For the 7T discovery dataset, we determined there was a Main Effect of Hemisphere (two-way ANOVA, *F*(1,388) = 24.09, *p* = 1.36 × 10^−6^) with the number of streamlines significantly higher in the left (PUL-MT, M = 16 +/− 19 and LGN-MT M = 13 +/− 14) compared to the right hemisphere (PUL-MT, M = 7 +/− 8 and LGN-MT M = 9 +/− 8). Similarly, for the 3T validation dataset, we observed a Main Effect of Hemisphere (two-way ANOVA, *F*(1,1528) = 273.67, *p =* 1.19 × 10^−56^) but also a Main Effect of Region (*F*(1,1528) = 34.84, *p* = 4.42 × 10^−9^) and an Interaction (*F*(1,1528) = 30.81, *p =* 3.35 × 10^−8^) with more streamlines connecting the left PUL-MT compared to the bilateral LGN-MT.

#### Apparent fibre density (AFD)

To investigate these results further, we used the outputs from SIFT2 (Smith et al., 2015) to determine the apparent fibre density (AFD) of the pathways connecting the ROIs (Figure 1C, bottom). The AFD expresses the ‘fibre weights’ of the reconstructed white matter tracts, with higher fibre densities corresponding to a greater number of axons comprising each tract. Note that we examined this measure only in individuals for whom we were able to reconstruct a tract. In the 7T discovery dataset, using a two-way ANOVA we observed a Main Effect of Hemisphere (*F*(1,357) = 17.68, *p* = 3.30 × 10^−5^) with higher fibre densities in the left compared to the right hemisphere. Post-hoc analyses revealed that the left PUL-MT tract had the highest fibre density (M = 0.10, SD = 0.13) and that the right PUL-MT (M = 0.05, SD = 0.06) and right LGN-MT (M = 0.05, SD = 0.05) had the lowest. In the 3T discovery dataset, using a two-way ANOVA, we observed a Main Effect of Hemisphere (*F*(1,1412) = 254.13, *p* = 9.78 × 10^−53^) and also a Main Effect of Region (*F*(1,1412) = 73.21, *p* = 2.98 × 10^−17^) and an Interaction (*F*(1,1412) = 53.42, *p* = 4.50 × 10^−13^). Post-hoc analyses revealed that, as in the 7T discovery dataset, the left PUL-MT tract had the highest fibre density (M = 0.88, SD = 0.94) and the right PUL-MT (M = 0.17, SD = 0.34) and right LGN-MT (M = 0.13, SD = 0.25) the lowest.

#### Pulvinar subnuclei connectivity

To examine the PUL connectivity in greater detail, we used additional masks to subdivide this ROI into the known anatomical subnuclei (that is, the superior, inferior, anterior, medial and lateral pulvinar subcompartments). We then used local tractography to reconstruct the PUL-MT white matter tracts that originated or terminated within these specific subnuclei (Figure 1D). Using this method for the 7T discovery dataset, we determined that the majority of the white matter fibres connecting the PUL to MT originated within the inferior PUL (mean number of streamlines in left hemisphere = 16, SD = 19 and right M = 9, SD = 9) and lateral PUL (left M = 6, SD = 8 and right M = 5, SD = 6). Similarly, in the 3T validation dataset, we found that the majority of the fibres connecting the PUL to MT originated again within the inferior PUL (left M = 94, SD = 121) and right M = 63, SD = 80) or the superior PUL (left M = 58, SD = 83 and right M = 42, SD = 58). Thus, the inferior pulvinar served as the main pathway through which the pulvinar connected to MT but note that all subnuclei had some level of connectivity (see Table 7 for further details).

### LGN-MT tracts are more myelinated

Once we reconstructed the subcortical pathways connecting MT to both PUL and LGN, we then asked which of these tracts was more highly myelinated and would give us a candidate for the pathway that transmits information most efficiently. For this, we extracted the T1w/T2w ratio as a proxy for axonal myelination (Ganzetti et al., 2014) that has been found to closely align with the underlying myelination when estimated indirectly (Barkovitch, 2005; Glasser & Van Essen, 2011). Using the local tractography results as masks, we extracted the T1w/T2w ratio per tract and participant and then obtained the mean value per tract (Figure 2). For the 7T discovery dataset, we observed a Main Effect of Region (two-way ANOVA, *F*(1,353) = 38.30, *p* = 1.68 × 10^−9^) with greater myelination of the LGN-MT pathways. Post-hoc statistics determined that the LGN-MT pathways (left M = 1.27, SD = 0.15 and right (M = 1.29, SD = 0.15) were more highly myelinated than the PUL-MT pathways (left M = 1.17, SD = 0.14 and right M = 1.21, SD = 0.13). Similarly, using the 3T validation dataset, we observed a Main Effect of Region (two-way ANOVA, *F*(1,1397) = 4.99, *p* = 0.0256) and Hemisphere (*F*(1,1397) = 30.12, *p* = 4.81 × 10^−08^). Post-hoc statistics revealed that the LGN-MT pathway was more highly myelinated (left M = 1.66, SD = 0.47 and right M = 1.55, SD = 0.41) than the PUL-MT pathways (left M = 1.62, SD = 0.47 and right M = 1.48, SD = 0.41), and more so in the left compared to the right hemisphere, respectively.

**Figure 2.**
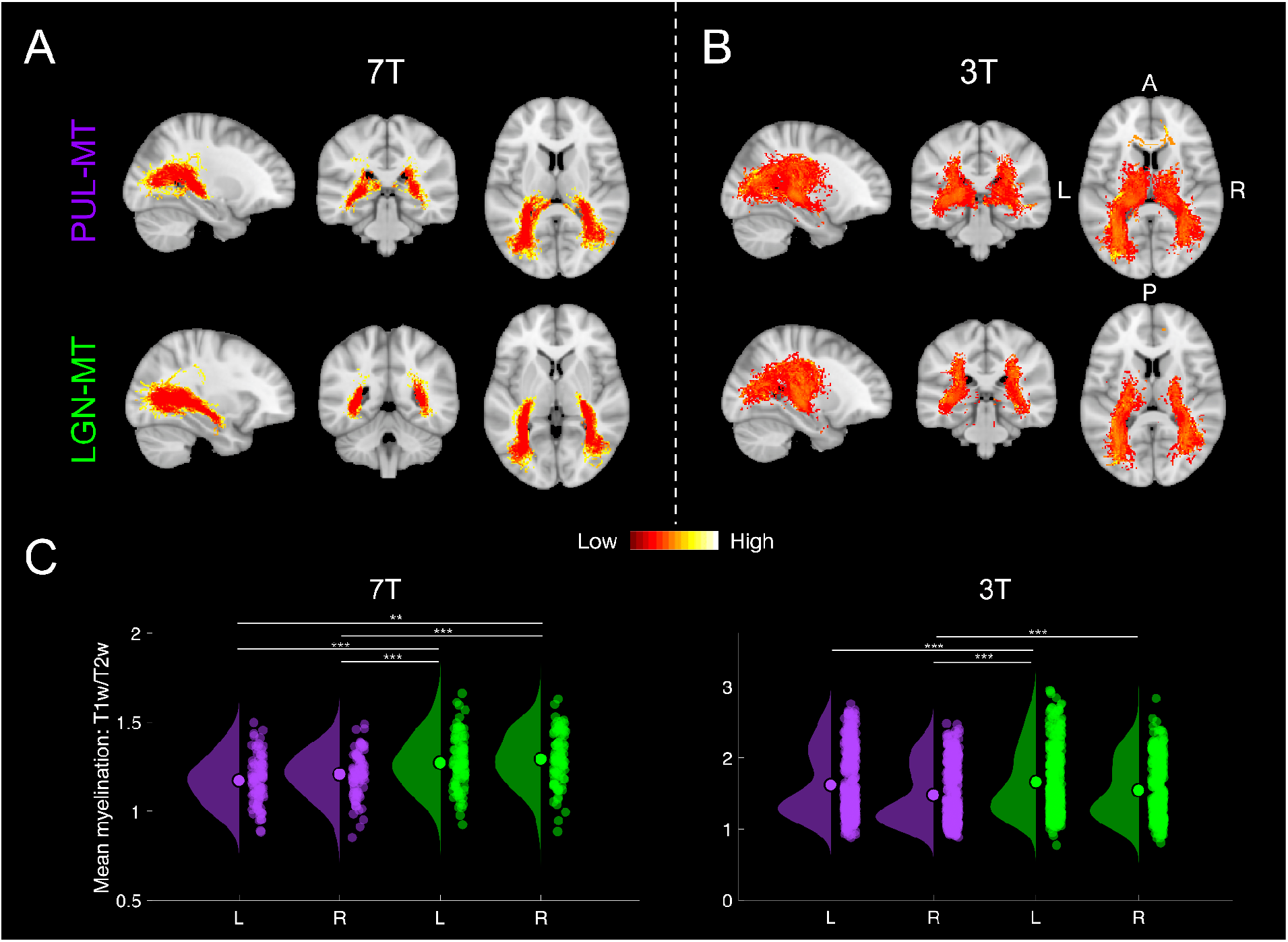
Mean myelination (T1w/T2w ratio) across the PUL-MT and LGN-MT pathways. **(A)** Visual representation of the mean myelination along the PUL-MT pathway (top) and LGN-MT pathways (bottom) for the 7T ‘discovery’ dataset and **(B)** for the 3T ‘validation’ dataset. For visualisation purposes, we combined the myelination values at every voxel for each participant and divided by the number of individuals with a tract within that voxel. **(C)** The mean myelination value (T1w/T2w ratio) for individual participants along the left and right PUL-MT (purple) and LGN-MT (green) pathway for the 7T (left) and 3T (right) datasets.

## DISCUSSION

In this study we used data (7T ‘discovery’ set, N = 98, and 3T ‘validation’ set, N = 381) from the HCP database to reconstruct subcortical visual pathways to MT from the PUL and the LGN. Overall, tracts in the left hemisphere tended to have a higher fibre count and greater fibre density (or AFD), suggesting left lateralised tracts might more efficiently transmit subcortical (arguably unconscious; Le Doux, 1996; for review see McFadyen et al., 2020) visual motion information compared to tracts in the right hemisphere. For the PUL subnuclei, we found that while all regions had some level of connectivity to MT, the greatest number of connections originated from the inferior pulvinar subnucleus, in keeping with previous findings (Bridge et al., 2016; Kastner et al., 2011; Bennaroch, 2015); and, again, more so within the left hemisphere. Interestingly, while the left PUL-MT pathway had the greatest number of individual streamlines and AFD, the LGN-MT pathways had the greatest mean level of myelination. Thus, while our findings showed a left hemispheric advantage, taken together, our results suggest subcortical motion signals might be carried *most* efficiently via the more highly myelinated LGN-MT pathways, and specifically along the left LGN-MT tract.

### Subcortical shortcuts for visual motion

Our study is, to the best of our knowledge, the first to reconstruct PUL-MT and LGN-MT pathways in a large sample of healthy individuals, with past studies having focused on non-human primates (e.g., Saalman & Kastner, 2011; Warner et al., 2014; Sincich et al., 2004) or small groups of patients with brain damage and age-matched controls (e.g., Ajina et al., 2015; Schmid et al., 2010; Barleben et al., 2015). In non-human primates, the LGN-MT pathway is thought to be predominantly magno- (Kaas & Baldwin, 2019) and koniocellular (Sincich et al., 2004), and responsible for the residual features underlying blindsight capabilities (that is, unconscious visual motion processing) after extensive damage to V1 (Sincich et al., 2004; Fries, 1981). In humans, the LGN is known to comprise parvo-, magno- and koniocellular layers from direct retinal inputs (Kastner, 2008) and the LGN-MT pathway to be critical for blindsight capabilities observed in adult individuals (as its absence predicts a lack of blindsight capabilities; Ajina et al., 2015; however, see Kinoshita et al., 2019 showing that the PUL pathway via the superior colliculus contributes to blindsight abilities in monkeys and Aghakhanyan et al., 2004 study showing strengthening of a PUL-MT pathway in an infant with blindsight caused by a gestational tumour). Interestingly, in healthy individuals, it has been suggested that direct LGN-MT connectivity might mediate rapid motion detection in everyday vision (e.g., >6°/s; ffytch et al., 1995; Gaglianese et al., 2012; but also see Schmid et al., 2010; Tamietto & Morrone, 2016) and deliver short-latency motion information to the dorsal stream (Ghodrati et al., 2017).

For the PUL-MT pathway, work in non-human primates has shown that the inferior and lateral PUL are reciprocally connected with MT (Standage & Benvento, 1983; Saalman & Kastner, 2011), suggesting this PUL-MT pathway may represent a motion-sensitive path that bypasses the LGN entirely. Indeed, in adult monkeys and humans, it has been suggested that early maturation of the visual motion pathways may rely on the PUL-MT pathways which are densely connected during neonatal development but later pruned as the LGN-V1-MT pathway dominates (Bourne & Morrone, 2017). Other research into early-life V1 lesions in marmosets suggest that some form of blindsight capabilities might also rely on a PUL-MT pathway being favoured over the LGN-V1 pathway as retinal input to LGN degenerates following such damage (Warner et al., 2014). Furthemore, in a recent review by Tamietto & Morone (2016), the authors suggested that the PUL-MT pathway (which also involves direct connections to the amygdala) might rapidly process motion signals with affective salience (such as those observed in affective blindsight; Tamietto et al., 2012; Rafal et al., 2015) while other studies suggest the PUL-MT pathway might mediate rapid motion detection and saliency processing (Furman et al., 2014) or maintain visual stability during rapid eye movements (Wurtz et al., 2008). Overall, our study provides evidence for transmission of visual motion signals via subcortical PUL-MT and LGN-MT pathways (with the most connections observed between the inferior PUL and MT; as observed in non-human primates); the functional significance of which is yet to be determined.

### Why two shortcuts?

But why would *two* subcortical pathways exist to carry (possibly unconscious) information about visual motion? And what sorts of visual information might they carry? Given the wide range of different types of visual motion that bombard us in everyday life, it seems reasonable that more than one subcortical visual motion pathway might exist to carry different kinds of specialised information. For example, some of the types of visual motion information might relate to: biological movement (of other organisms in space), object movement and tracking (of inanimate items in space), optic flow caused by the movement of our own body in space (e.g., to use for balance and the detection of our own movement direction) and the movement of the body and facial expressions in social contexts; in addition to information relating to the speed and direction of motion. Furthermore, while the traditional model of visual processing is that information flows along either a dorsal (motion, ‘where pathway’) or ventral (object, ‘what pathway’)’ route (Baizer et al., 1991), recent work suggests that visual motion might play a much more fundamental role in visual perception than first believed and that its function may extend past that of the dorsal visual pathway and further into the visual processing hierarchy (Gilaie-Dotan, 2016). Thus, the notion of two subcortical visual motion pathways seems necessary and practical to support this vast array of functions.

Another potential reason for two subcortical visual motion pathways may be to transmit the different information carried by the magno-, parvo- or koniocellular pathways. Indeed, early evidence in non-human primates suggests the LGN-MT pathway may be primarily magno- and koniocellular (Kaas & Baldwin, 2019; Sincich et al., 2004; however, note the case for parvocellular inputs; Anderson et al., 1995; Nassi et al., 2012). On the contrary, work examining the PUL-MT tract in monkeys suggests predominance of parvo- and koniocellular types (Huo et al., 2019; Wen et al., 2021). And while magnocellular neurons are known to preferentially respond to motion, (Butler et al., 2001) colour information (red-green carried in parvocellular layers and blue-yellow in koniocellular layers; Floyd et al., 2004; Yoonesi & Yoonesi, 2011; Ahmadi et al., 2015) can also influence motion perception by providing clues on form, apparent movement and depth (Ffytche et al., 1995b; Dobkins, 2000). Thus, both magno- and parvocellular layers might be necessary for the perception of motion (De Yoe & Van Essen, 1988; Lennie et al., 1990; Anderson et al., 1995; De Valois et al., 2000), while it is still unclear how the koniocellular layers contribute to motion perception. Thus, future studies that exploit the different responses of these cell types (specifically, their relation to visual motion perception) will be needed to elucidate the functions of these two subcortical pathways.

### Left hemispheric dominance of both subcortical visual motion pathways

Another interesting question that arises from the findings of this study is: why do both the PUL-MT and LGN-MT subcortical visual motion pathways exhibit left hemispheric dominance (that is, a presumed increase in processing capacity based on greater streamline counts and higher AFD)? Indeed, left and right visual field and hemispheric asymmetries in motion perception have been reported in a small number of studies (see below), however, the current neuroimaging study is the first in such a large sample of healthy individuals. While we did not initially predict this asymmetry, there is evidence to suggest a right visual-field left-hemispheric (RVF-LH) advantage for visual motion perception with the left hemisphere proposed to have finer temporal processing resolution, potentially attributable to the reading direction of western societies (Nicholls & Atkinson, 1993; Okubo & Nicholls, 2005; 2008; Levy et al., 2010). For example, an early study by Morikawa & McBeath (1992) found a robust bias for leftward movement in western readers that was not observed in bilingual individuals whose first language read from right to left. Similarly, Okubo & Nicholls (2005) used nonverbal stimuli presented either to the left or right visual field and determined there was a RVF-LH advantage for short duration stimuli presented on green (but not red) backgrounds. The authors suggested these findings could be attributed to the magnocellular pathway (as red wavelengths disrupt the functioning of these cells) driving the left hemisphere advantage at short temporal scales. More recently, Rima and colleagues (2020) showed a significant RVF-LH advantage for detecting visual events in the right hemifield which they also attributed to left hemispheric dominant reading habits.

On the other hand, some studies have reported inconclusive or contradictory evidence regarding the hemispheric lateralisation associated with visual motion processing. In one study with a small sample (N = 6) of healthy individuals, the authors found that the onset of visual motion evoked potentials in adults (specifically the N2-component) were driven by the right or left hemisphere (not predicted by handedness) regardless of whether the stimuli were presented in the left- or right-visual hemifield (Hollants-Gilhuijs et al., 2000). Conversely, Caso & Spinelli (1988) presented horizontally moving motion stimuli within the left or right visual field (at 4 deg eccentricity) and found that right-handed participants had a left visual field sensitivity advantage while the opposite was true for left-handed participants. Note that the ‘handedness’ hypotheses do not align with our findings as, in our study, the majority of participants (91% 7T and 93% 3T) were right-handed and yet we observed LH strengthening. Such LH increases in AFD and myelin are presumably related to RVF advantages, although we lack the behavioural data in this study to make a direct link between the two. Finally, Strong and colleagues (2019) found greater deficits in motion perception when TMS was applied to the right compared to the left hemisphere and suggested a greater role for the right hemisphere in full-field motion processing. Thus, while there is evidence in support of our findings of a RVF-LH motion processing advantage, our results do not align with contradictory findings of right hemispheric advantages based on handedness, and it is unclear whether these effects are driven by the cortical or subcortical pathways. Future work will be needed to determine whether our results are observed in other samples of healthy individuals and to determine how these findings influence visual motion processing.

### Greater number of denser fibre bundles in the left PUL tracts are less myelinated than the LGN pathways

While the left PUL-MT tract had the most connecting white matter fibre bundles that comprised a greater number of axons, these tracts were not as highly myelinated as the LGN-MT pathways. This finding is somewhat surprising. As axon diameter and myelination are positively correlated and represent the most important factors influencing the speed of neural transmission (Caminiti et al., 2012; Friedrich et al., 2020; Bechler et al., 2015), one might expect the most efficient pathways to contain more streamlines, greater fibre density and the highest level of myelination. Interestingly, however, recent work suggests that axonal myelination is *both* size- and experiencedependent (Stadelmann et al., 2019; Bechler et al., 2015; Mount & Monje, 2017), with larger diameter axons (> 1um) tending to be myelinated earlier and more completely (Stadelmann et al., 2019) and with activity-dependent models of neural plasticity also regulating axonal myelination (Bechler et al., 2015; Mount & Monje, 2017). Indeed, a recent review by Mount & Monje (2017) highlights how ‘adaptive myelination’ in sensory processing circuits (as well as other cortical regions) may allow for the subtle tuning of perceptual features, which would be especially important for temporally specific processing early in the sensory hierarchy. Based on these findings, we suggest that the denser left hemispheric pathways might be more highly myelinated based on their size, and the LGN-MT pathways myelinated even further based on these pathways being used more often. That is, if the adaptive myelination theory is correct in this situation, the LGN-MT pathway may be more heavily relied upon than the PUL-MT pathways and, therefore, more highly myelinated despite being less dense. Exciting future research into the adaptive myelination hypothesis, specifically, within these pathways, will shed light on these findings more deeply.

### Limitations and future directions

An intrinsic limitation of using MRI-based non-invasive DTI to reconstruct white-matter pathways is that this methodology uses external measurement of the apparent anisotropic movement (or diffusion) of hydrogen protons of water molecules (Merboldt et al., 1985; Taylor & Bushell, 1985). As such, DTI-based fibre tractography represents an indirect estimation of the white-matter tracts of the brain, which can be susceptible to false positives and negatives (Maier-Hein et al., 2017; de Reus & van den Heuvel, 2013), especially in areas of crossing fibres (Raffelt et al., 2017; Dhollander et al., 2021). These limitations are slowly being aided by the advancement of DTI analysis software such as those used in the current study and while probabilistic tractography can lead to more false positives than deterministic tractography, it is nevertheless advantageous when examining regions with a high density of crossing or fanning fibres (Zalesky et al., 2016). Another limitation of the current study is the use of cortical atlases set to identify the ROIs, instead of using an individual-specific approach, which is unfeasible here with such large samples. However, we can be reasonably certain that the use of our ROIs encapsulated the majority of brain regions housing the PUL, LGN and MT.

While the inherent limitations of using DTI for white-matter tract reconstruction can only be overcome with advancements in technology and modelling approaches, there are a number of ways in which the current study can be extended in the future. Specifically, the next steps for investigating these subcortical pathways for visual motion perception will be to design an fMRI task that, combined with DTI, will allow for a more in-depth investigation into the functional role of the PUL-MT and LGN-MT pathways. For this, one will need: (1) subject-specific ROI identification, (2) an experimental paradigm that manipulates different types of visual motion and (3) a means of investigating the effective connectivity between the ROIs (for example using Dynamic Causal Modelling, Friston et al., 2003). Firstly, it will be necessary to use a range of visual motion stimuli for retinotopic mapping of area MT in individual participants, and a separate set of auxiliary tasks that allow for individual-specific identification of the PUL and LGN regions. Secondly, a task that utilises different kinds of visual motion stimuli and which manipulates the response functions of the mango-, parvo- and koniocellular pathways will be necessary to pinpoint the roles for two subcortical visual motion pathways. With further investigation, the behavioural function of a dual PUL-MT and LGN-MT subcortical pathway for visual motion information will be elucidated.

## CONCLUSIONS

In this study, we used DTI-based local probabilistic tractography to reconstruct two subcortical visual motion pathways connecting PUL-MT and LGN-MT in a large sample of healthy individuals using 7T ‘discovery’ and 3T ‘validation’ datasets. We found a left hemispheric lateralisation of these pathways as evidenced by a greater number of streamlines with greater fibre densities and myelination. We suggest this lateralisation may emerge with western reading habits that lead to advantageous processing of motion in the right visual hemifield, however, further work will be needed to confirm the nature of this lateralisation. We also show that pathways connecting LGN-MT were more highly myelinated than PUL-MT pathways. We suggest that, despite a greater number of fibres with higher density within the left PUL-MT tract, the greater level of myelination in the LGN-MT tracts may be due to ‘adaptive myelination’ caused by more frequent use that leads to faster signal transduction. Further research will be needed to determine the types of visual motion signals that are carried by both the PUL-MT and LGN-MT pathways and what kinds of functions they subserve.

